# Variations of gut microbiome profile under different storage conditions and preservation periods: A multi-dimensional evaluation

**DOI:** 10.1101/752584

**Authors:** Junli Ma, Lili Sheng, Chuchu Xi, Yu Gu, Ying Hong, Ningning Zheng, Linlin Chen, Gaosong Wu, Yue Li, Juan Yan, Ruiting Han, Bingbing Li, Huihui Qiu, Jing Zhong, Wei Jia, Houkai Li

## Abstract

Gut dysbiosis contributes to the development of various human diseases. There are thousands of publications per year for investigating the role of gut microbiota in development of various diseases. However, emerging evidence has indicated data inconsistency between different studies frequently, but gained very little attention by scientists. There are many factors that can cause data variation and inconsistency during the process of microbiota study, in particular, sample storage conditions and subsequent sequencing process. Here, we systemically evaluated the impacts of six fecal sample storage conditions (including −80 °C, −80 °C with 70% ethanol (ET_-80 °C), 4°C with 70% ethanol (ET_4°C), and three commercial storage reagents including OMNIgene•GUT OMR-200 (GT), MGIEasy (MGIE), and Longsee (LS)), storage periods (1, 2 weeks or 6 months), and sequencing platform on gut microbiome profile using 16S rRNA gene sequencing. Our results suggested that −80°C is acceptable for fecal sample storage, and the addition of 70% ethanol offers some benefits. Meanwhile, we found that samples in ET_4 °Cand GT reagents are comparable, both introduced multi-dimensional variations. The use of MGIE resulted in the least alteration, while the greatest changes were observed in samples stored in LS reagents during the whole experiment. Finally, we also confirmed that variations caused by storage condition were larger than that of storage time and sequencing platform.

**IMPORTANCE:** In the current study, we performed a multi-dimensional evaluation on the variations introduced by types of storage conditions, preservation period and sequencing platform on the basis of data acquired from 16S rRNA gene sequencing. The efficacy of preservation methods was comprehensively evaluated by DNA yield and quality, α and β diversity, relative abundance of the dominant bacteria and functional bacteria associated with SCFAs producing and BAs metabolism. Our results confirmed that variations introduced by storage condition were larger than that of storage periods and sequencing platform. Collectively, our study provided a comprehensive view to the impacts of storage conditions, storage times, and sequencing platform on gut microbial profile.

## INTRODUCTION

The mammalian gastrointestinal tract is the main site for commensal bacteria (1, 2), which contains at least 100-times as many genes as host genome (3). In recent years, the passion on gut microbiota-related research is overwhelming due to the involvement of gut dysbiosis in development of various human diseases including obesity, diabetes mellitus, nonalcoholic fatty liver diseases, cardiovascular disease, and even cancers (4-7). Emerging high-throughput sequencing technologies including 16S rRNA gene and metagenomics lay the solid foundation for investigating the role of gut microbiota in human diseases (8, 9).

There are over thousands of microbial-related publications per year mainly by using 16S rRNA gene sequencing technology. Unfortunately, gut microbiome data usually show dramatic variations and poor consistency between different reports (10-12). There are various ways for introducing disturbance on the diversity and composition of microbiome during the whole experimental process including sample collection, transportation, storage, DNA extraction, sequencing, and biometric analysis (13-15). Data derived from 103 fecal samples of two infant cohorts show that fecal microbial structure changes significantly during ambient temperature storage after 2 days, so immediate freezing at −80 °C or DNA extracted within 2 days is suggested if samples were stored at room temperature (16). However, another study reveals that fecal samples stored at room temperature beyond 15 minutes result in obvious variation in bacterial taxa, whereas usage of microbial nucleic acid stabilizer RNAlater also causes dramatic reduction in DNA yields and bacterial taxa(17). Generally, immediate freezing at −80 °C is supposed to be the gold standard for most biological samples including feces for microbiome study (18, 19). However, obvious alteration in microbial community is also observed in samples frozen at −80 °C compared to that of fresh samples, suggesting that −80 °C might not be the most optimal method for fecal sample storage (20). Moreover, immediate freezing at −80 °C is usually unfeasible in many cases for field studies, or at the circumstance where the fecal samples are collected at home by patients themselves. In these cases, samples are apt to be exposed at ambient temperature during collection and transportation before DNA extraction.

Currently, various commercial and experimental preservation reagents for fecal sample have been developed or trialed such as the OMNIgene Gut kit, RNAlater, FTA cards, 70% or 95% ethanol, 50:50 glycerol:PBS, NOBP-based reagent and so on (21-23). Although some comparisons between these preservation reagents have been performed, inconsistent or even contradictory results are observed. Song et al. find that the preserving effect of 95% ethanol is comparable with that of FTA cards or OMNIgene Gut kit at ambient temperature, but strongly caution against the use of 70% ethanol for fecal sample preservation (22). Contrarily, Horng et al. report that the microbial composition of samples preserved with 70% ethanol or RNAlater closely resembles that of fresh samples, and they suggest that 70% ethanol is the best method for preserving canine fecal samples (20). Similarly, contradictory results are also observed in the use of RNAlater. Study shows that fecal samples preserved with RNAlater closely resembles that of fresh samples (20), but contrary results are observed in another study, in which RNAlater-preserved samples have the least similarity in microbial composition and abundance with fresh samples (24). In addition, significant differences are also observed in microbial profile of identical samples that processed and sequenced at two research centers (25, 26). Collectively, these contradictory or inconsistent results highlight the significance and urgency for a further systemic evaluation on the variations of gut microbiome profile that could be introduced during the process of sample preparation such as different fecal storage conditions, period of preservation and even different sequencing platforms.

In the current study, we systemically evaluate the variations of gut microbiome profile with freshly collected and homogenized feces sample from rats that was allocated into various replicates for observing the impacts of 3 commonly used storage conditions including −80 °C, addition of 70% ethanol at either −80 °C (ET_-80 °C) or 4 °C (ET_4 °C), and 3 commercial stabilizers including OMNIgene•GUT OMR-200 (GT) and MGIEasy (MGIE) at ambient temperature, and Longsee (LS) at 4 °C according to their instructions. Bacterial genomic DNA was extracted with same protocol at the end of the 1^st^ week, 2^nd^ week, and 6^st^ month respectively, as well as the fresh samples. The study design was described in Fig. 1. All of the samples were subjected for gut microbiome profiling by using 16S rRNA gene sequencing. Our results demonstrated that fecal storage conditions did have dramatic impacts on gut microbiome profile including DNA quality, α or β diversity, relative abundance of functional bacteria involved in SCFA production and bile acid metabolism. We confirmed that −80 °C was acceptable for fecal sample storage, and the addition of 70% ethanol was plus for maintaining the original community composition of the dominant phyla. Meanwhile, we found that gut microbiome profiles of samples stored in ET_4°C and GT reagents were comparable regarding to their impacts on community α or β diversity, and the relative abundance of the dominant and functional bacteria. Our results showed that gut microbiome profile of samples stored in MGIE reagents was the closest to that of fresh samples, while LS reagents introduced the most obvious variation. Finally, we also confirmed that variations caused by ways of storage condition were larger than that of storage periods and sequencing platform.

**FIG 1.**
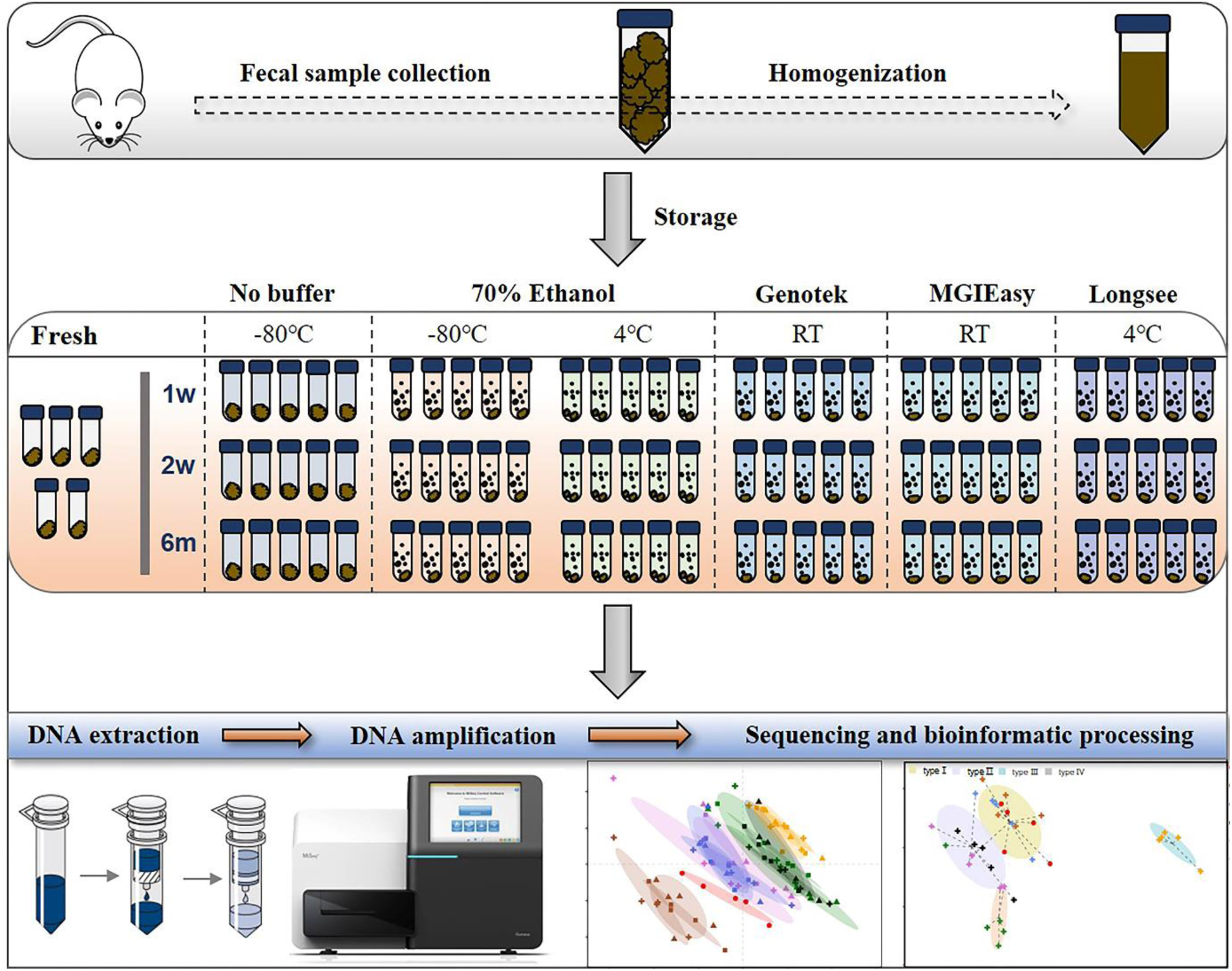
Diagram of experimental design. Fecal samples from 10 rats were quickly collected into sterile 50 ml tubes, homogenized, and then aliquoted. Next, these aliquots were treated with a range of preservatives. Bacterial DNA were extracted at 4 time points: on the day of sampling (“Fresh”) as well as after 1 week, 2 weeks, and 6 months of storage, followed by DNA extraction and 16S rRNA gene sequencing.

## RESULTS

### Do different storage conditions lead to variation in DNA yield or quality?

Qualified DNA is the basis for microbial study, which varies among different DNA extraction protocols (14). To determine the impact of fecal storage conditions on bacterial genomic DNA quality, we evaluated the concentration and purity of DNA extracted with same protocol from samples under different storage conditions. Despite obvious fluctuations, our results showed that DNA concentrations were comparable among most samples, except for the relatively higher in MGIE and lower DNA concentrations in GT and LS reagents at the 3 time points of storage (Fig. 2a). Meanwhile, DNA quality was evaluated with the absorption ratio of 260/280 nm. We found that most samples showed satisfactory value in 260/280 from 1.8 to 2.0, except for samples stored in LS reagents with relatively lower 260/280 value. Notably, DNA concentration of samples stored in GT reagents showed obvious reduction with time (Fig. 2b). Taken together, our current data suggested that most storage conditions had minor and acceptable impacts on DNA yields or quality, whereas samples in LS reagents showed relatively lower yields and quality. Preservation of samples in GT reagents might result in reduction of DNA yield time-dependently.

**FIG 2.**
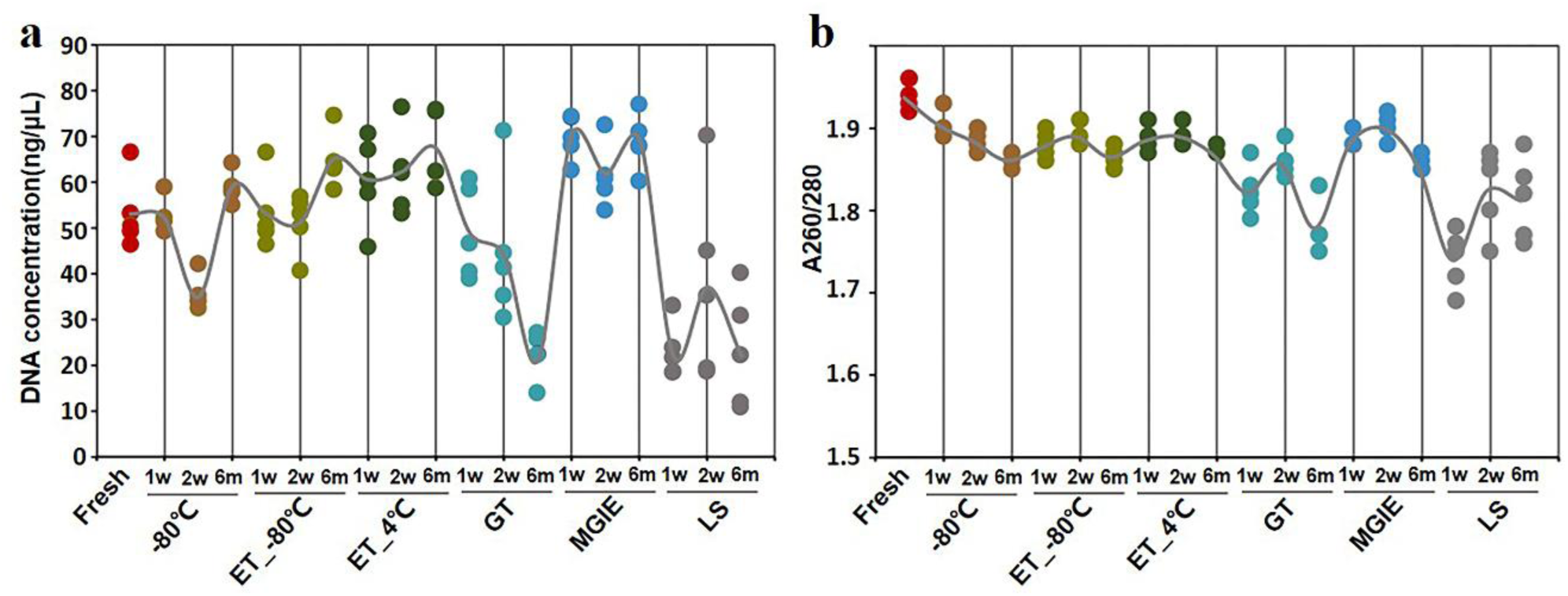
DNA quality under different storage conditions. (a-b) The concentration and purity of DNA extracted from fecal samples under different storage conditions. Curves shown were the average of five samples.

### Do different storage conditions affect bacterial diversity?

Bacterial diversity is the main character for investigating the role of gut microbiota in disease. First, we compared the bacterial α diversity with Shannon and Simpson indices, as well as Chao1 and Shannon’s evenness estimators under different storage conditions. Our results showed that the bacterial α diversity was differently altered in samples stored at - 80 °C for 1 or 2 weeks, and 6 months, ET_-80 °C and ET_4 °C for 1 week, as well as GT and MGIE reagents for 1 week, characterizing as significant lower Shannon and higher Simpson diversity indices compared with fresh samples. Notably, samples stored in LS reagents showed higher Shannon and lower Simpson diversity indices at the 3 time points compared with fresh samples (Fig. 3a-b). Although the operational taxonomic unit (OTU) richness did not differ significantly among storage conditions, considerable variation in evenness was observed. In addition, samples in LS reagents showed the most significant variation in α diversity. (Fig. 3c-d).

**FIG 3.**
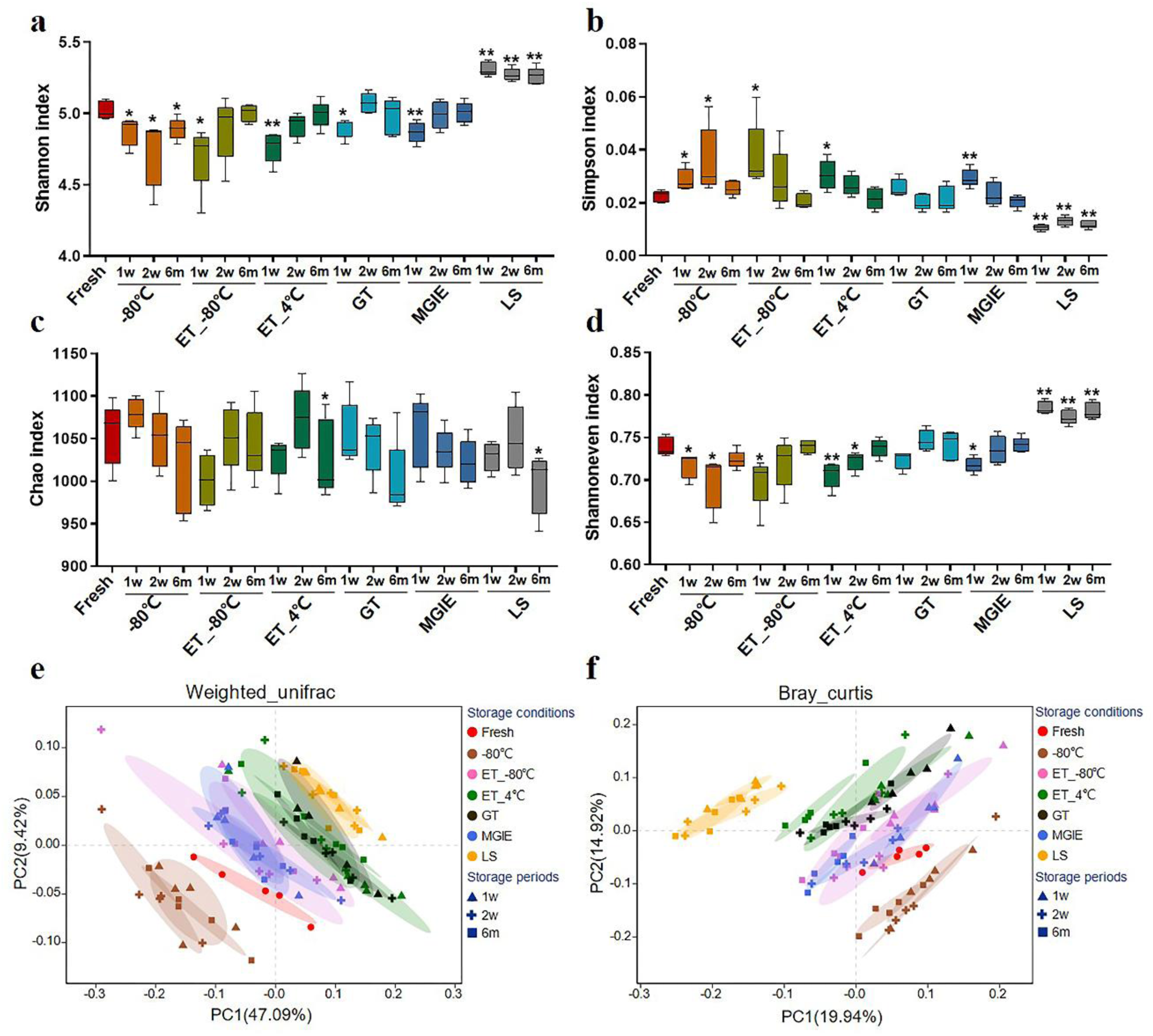
The effects of preservation methods on community structure. Community α diversity was measured by (a) Shannon diversity index and (b) Simpson diversity index. Community richness and evenness were evaluated by (c) Chao1 estimator and (d) Shannon index-based measure of evenness. Community β diversity was measured by PCoA plots based on (e) Weighted UniFrac and (f) Bray-Curtis. * *p* < 0.05, \*\**p* < 0.01, compared with fresh group.

Next, Principal Coordinate Analysis (PCoA) based on Weighted UniFrac and Bray_curtis were performed, which showed obvious variation in microbial community under different storage conditions. Generally, samples from the same storage condition for different periods were clustered together compared to storage conditions, suggesting that impacts of storage conditions are larger than storage periods. Specifically, samples stored at −80 °C, ET_-80 °C and MGIE reagents were clustered closely to fresh samples, and followed by those in GT or ET_4 °C, while samples in LS showed the worst clustering with fresh samples (Fig. 3e-f). Collectively, these results suggested that storage conditions had dramatic impact on α or β diversity of gut microbiota to different extent compared with their fresh control, and the impact of storage condition superseded that of storage periods.

### How do different storage conditions affect the abundance of dominant bacteria?

Given the observed impacts of storage conditions on bacterial α or β diversity, we further investigated the variations in the relative abundance of the dominant bacteria by comparing the top 60 OTUs from Firmicutes (30 OTUs), Bacteroidets (28 OTUs) and Proteobacteria (2 OTUs) phyla under different storage conditions which accounted for about 65% of coverage. As the heatmap showed in Fig. 4, we found that different storage conditions resulted in dramatic changes to different extent in abundance of most OTUs compared to fresh samples. By contrast, we found that the numbers of altered OTUs in samples at −80 °C, ET_-80 °C, and MGIE reagents were relatively smaller than those in ET_4 °C, GT or LS reagents. Interestingly, the majority of altered OTUs in samples stored at −80 °C showed decreased abundance, while addition of 70% ethanol could balance the ratio of up- or down-regulated bacteria at −80 °C. It was also notable that the OTUs abundance under different storage conditions were changed either time-dependent or –independently. For example, the samples stored at −80 °C, ET_4 °C and MGIE resulted in time-dependent increase in number of up-regulated bacteria, however, the number of down-regulated bacteria was much random under most conditions. In addition, much more universal alteration was observed in OTUs belonging to Firmicutes phylum under different storage conditions than that of Bacteriodetes.

**FIG 4.**
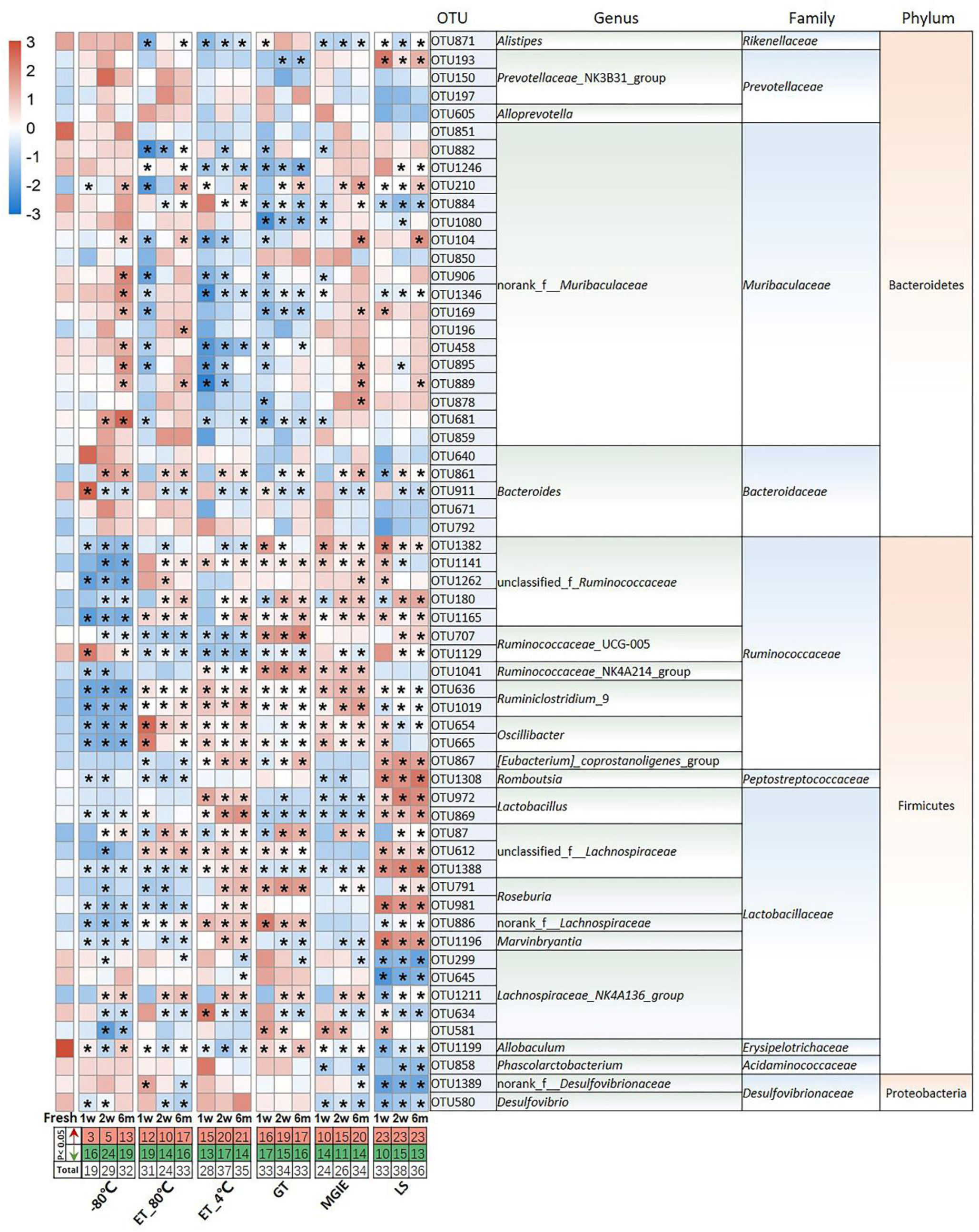
The impact of preservation methods on the top 60 OTUs. Heatmaps shown was the differences in relative abundance among groups of the top 60 OTUs, which accounted for about 65% of coverage, as well as their taxa information including genus, family, and phylum. The red and green entries indicate the number of OUT that were significantly more or less abundant under different storage conditions relative to fresh samples respectively. * *p* < 0.05, compared with fresh group.

We next evaluated the change in relative abundance of the 3 dominant phyla, Firmicutes, Bacteroidets, and Proteobacteria, as well as genera under different storage conditions by comparing with fresh controls individually. In comparison with fresh controls, −80 °C storage resulted in obvious increase in Bacteroides and decrease in Firmicutes time-dependently, but not Proteobacteria. Interestingly, addition of 70% ethanol at −80 °C (ET_-80 °C), but not 4 °C (ET_4°C) storage showed benefit for keeping the abundance of both Firmicutes and Bacteroidets close to fresh control at the 3 time points, except for Proteobacteria. Samples stored in either GT or LS reagents resulted in obvious increase in Firmicutes and decrease in Bacteroidetes time-dependently, as well as decreased abundance of Proteobacteria only in LS reagents. Notably, the relative abundance of the 3 dominant phyla was well maintained at the 3 time-points in MGIE reagents storage compared to fresh control (Fig. 5a).

**FIG 5.**
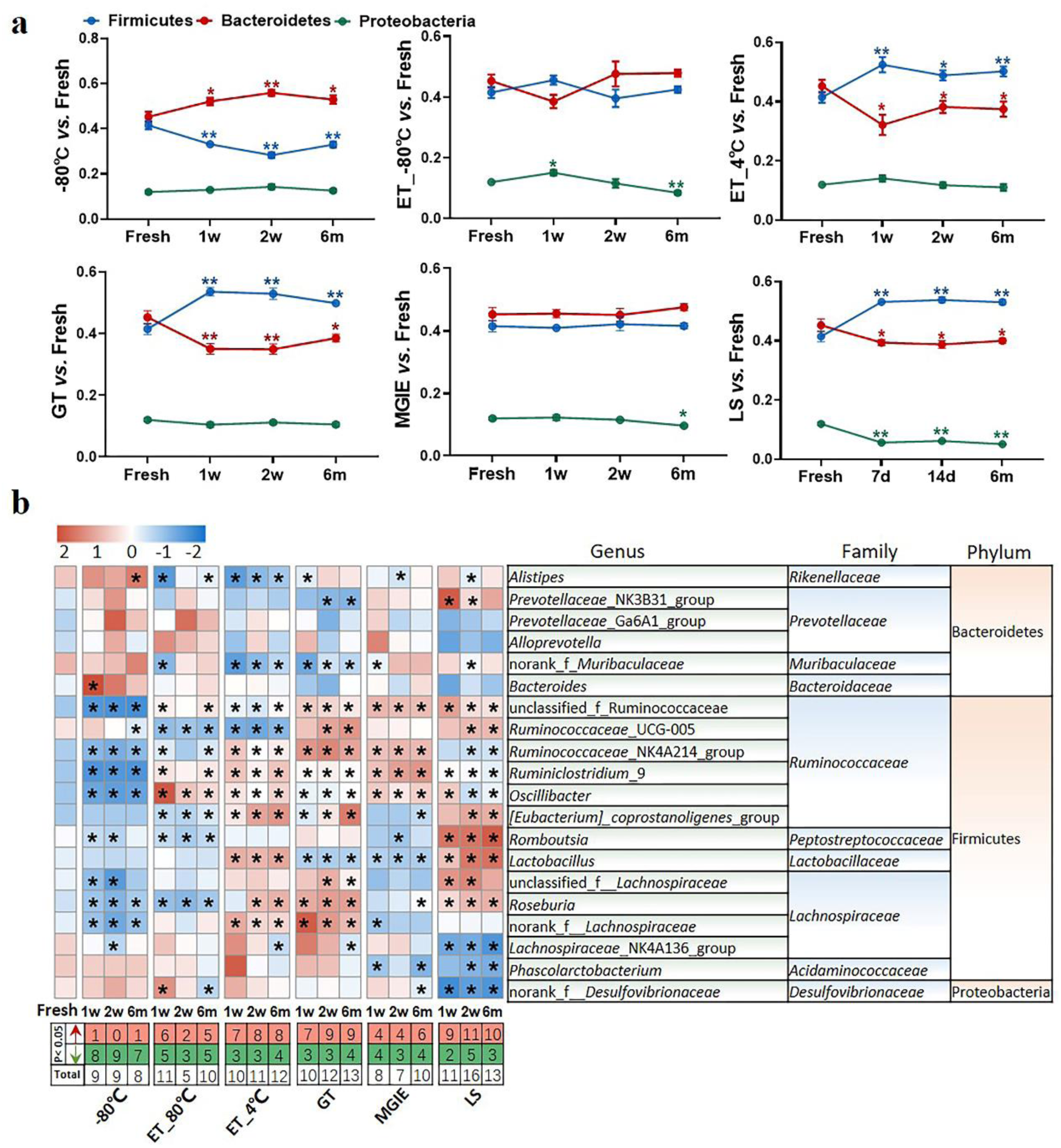
The effect of preservation methods on the dominant bacteria. (a) The effect of storage conditions on the relative abundance of the dominant phyla. (b) The heatmap shown was the differences in relative abundance among groups of the top 20 genera, which accounted for about 85% of coverage, as well as their taxa information including family and phylum. The red and green entries indicate the number of genus that were significantly more or less abundant under different storage conditions relative to fresh samples respectively. *p* < 0.05, \*\**p* < 0.01, compared with fresh group.

The imbalanced ratio of Firmicutes to Bacteroidetes (F/B) was frequently observed in many diseases (27-30), therefore, the evaluation of F/B ratio is of significance for investigating the role of gut microbiota in disease development or drug efficacy. Our results showed that F/B ratios of samples in ET_-80 °C and MGIE reagents were of little difference to that of fresh samples, whereas increased F/B ratios were observed in ET_4 °C, GT and LS reagents, but decreased in −80 °C (Fig. S1a). Consistent with previous results at phylum level, further analysis of the top 20 genera (85% of coverage) showed that samples in - 80 °C, ET_-80 °C, and MGIE reagents introduced the minimum biases, followed by ET_4 °C and GT reagents, whereas LS reagents caused the most obvious variation with up to 40 altered genera in total. Interestingly, we found that the majority of altered genera in −80 °C are decreased abundance at 3 time points, while addition of 70% ethanol at −80 °C made the number of increased or decreased genera relatively balanced. On the contrary, alterations introduced by ET_4 °C, GT and LS reagents were mainly increased. It was found that most of the variable genera are belonging to Gram-positive *Ruminococcaceae*, such as *Ruminococcaceae_*UCG-005, *Ruminiclostridium*_9, and *Oscillibacter*, whereas bacteria in genera of *Prevotellaceae_*NK3B31_group, *Alloprevotella* and *Bacteroides* were of minor change under different conditions, which are belonging to Gram-negative *Prevotellaceae*, and *Bacteroidaceae.* Nevertheless, we did not observe the time-dependent impacts of storage periods on number of altered genera under all of the storage conditions (Fig. 5b). In addition, correlation analysis between observed conditions and fresh samples was performed with Spearman’s correlation coefficient (SCC). Our results showed that samples in −80 °C and ET_-80 °C have the highest correlation with fresh samples ranging from 0.85 to 0.9, followed by ET_4 °C, GT, and MGIE reagents ranging from 0.75 to 0.85. Samples stored in LS reagents exhibited the worst similarity to fresh samples, in which the SCC was less than 0.7(Fig. S1b). Collectively, our results indicated that samples in ET_-80 °C and MGIE reagents kept the dominant phyla relatively stable including Firmicutes, Bacteroidetes, and Proteobacteria at 3 time points. Samples in LS reagents showed relatively higher variations at all levels. Moreover, the impacts of short- or long-term storage were not very significant under the same condition.

### Do different storage conditions alter the relative abundance of functional bacteria?

Increasing evidence have confirmed that gut microbiota play critical roles in maintaining human health or disease development by producing microbial metabolites like bile acids or SCFAs that serve as signaling factors or energy substrates (31). Therefore, we evaluated the relative abundance ratio (stored samples / fresh samples) of bacteria at genus level at 3 time points that are reported to be involved in SCFA-producing including genera of *Lachnospiraceae_*NK4A136_group, *Roseburia, Prevotella_9*, and *Blautia*, as well as that are involved in both SCFA-producing and bile acid metabolism including *Bacteroides, Lactobacillus* and *Ruminococcus_1* (32, 33). First of all, we found that even −80 °C storage from the sample collection caused some changes of these bacteria at genus level such as obvious reduction of *Lachnospiraceae_*NK4A136_group, *Roseburia* and *Blautia*, and increasing of *Prevotella_9* and *Bacteroides*, and no time-dependent impacts were observed. Addition of 70% ethanol at −80 °C produced similar effect with −80 °C, except for some benefits in *Prevotella_9, Bacteroides*, and *Lachnospiraceae_*NK4A136_group which were closer to fresh samples. However, storage at ET_4 °C caused dramatic variations in most of these bacteria at a time-dependent way, except for *Bacteroides.* There were also obvious variations in abundance ratio of these bacteria in the 3 commercial stabilizers. Interestingly, the change trends were much more similar between GT and MGIE such as bacteria in *Blautia, Lactobacillus* and *Ruminococcus_1*, while samples stored in LS reagents showed dramatic differences in majority of the observed bacteria, especially the unique change in *Lachnospiraceae_*NK4A136_group, *Blautia, Lactobacillus* and *Ruminococcus_1* (Fig. 6 a-b). We also found that *Bacteroides* and *Lachnospiraceae_*NK4A136_group were relatively stable under these storage conditions. Thus, our data suggested that different storage conditions could cause diversified fluctuations in the relative abundance of functional bacteria in a time-dependent or –independent way.

**FIG 6.**
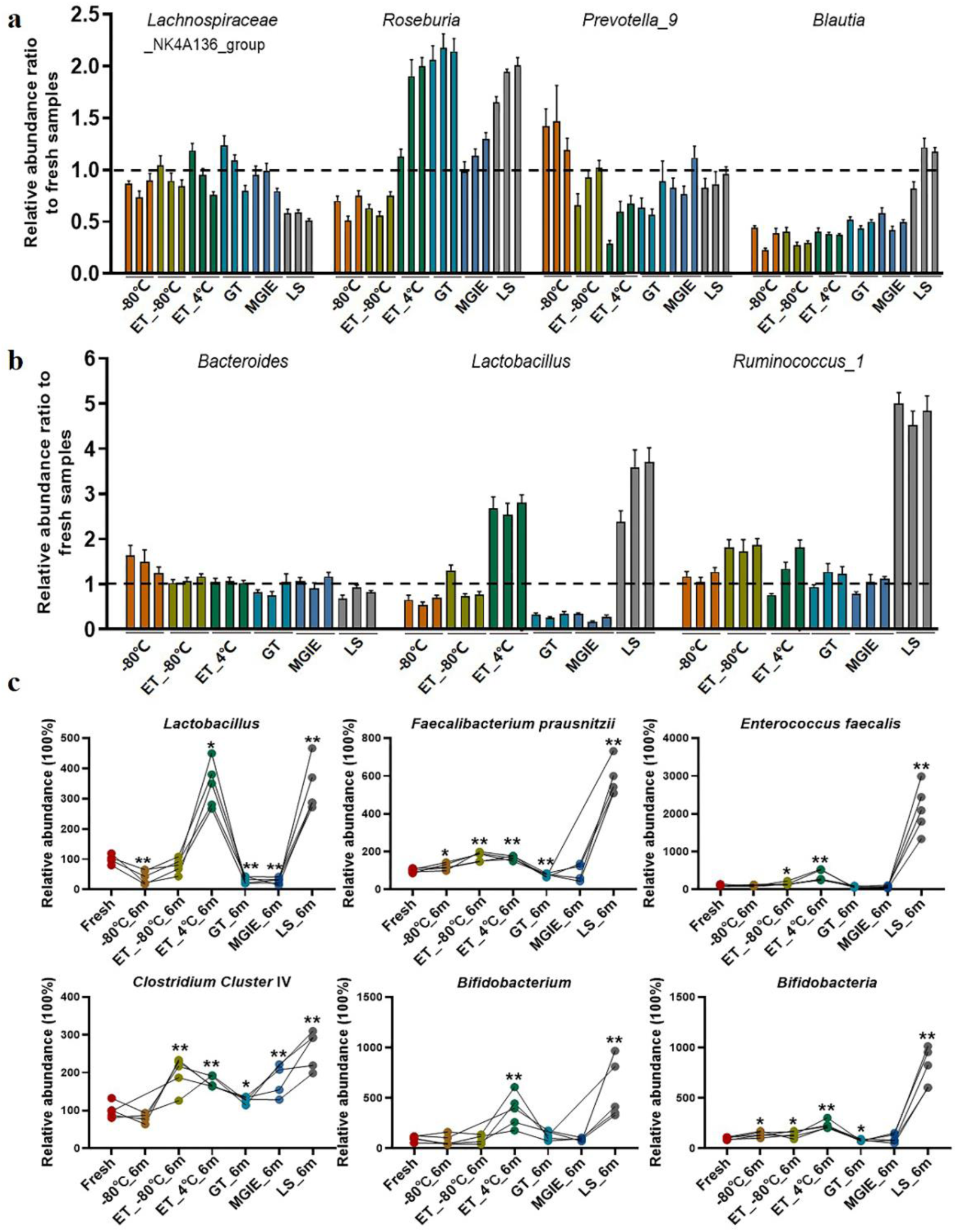
Relative abundance ratio of bacteria involved in SCFA-producing and bile acid metabolism on genus levels. (a-b) Relative abundance ratio (storage samples / fresh samples) of bacteria involved in SCFA-producing on genus levels, and three columns with the same color represent 1 week, 2 weeks, and 6 months from left to right respectively. (c) Qualification of the functional bacteria relative to fresh samples using QPCR. *p* < 0.05, \*\**p* < 0.01, compared with fresh group.

In addition, to further confirm the effect of storage conditions on the functional bacteria, we qualified the abundance of bacteria involved in BAs metabolism and SCFAs metabolism (*Lactobacillus, Faecalibacterium prausnitzii, Enterococcus faecalis, Clostridium Cluster* IV, *Bifidobacterium, Bifidobacteria*) with qPCR in samples stored for 6 months. We found that different storage conditions resulted in dramatic alterations to different extent in abundance of most bacteria compared to fresh samples. Detailly, −80 °C storage caused alterations in *Lactobacillus, Faecalibacterium prausnitzii*, and *Bifidobacteria*. Addition of 70% ethanol at −80 °C produced similar effect with −80 °C in *Faecalibacterium prausnitzii* and *Bifidobacteria*, and provided the benefit in *Lactobacillus*, while caused the increased abundance of *Enterococcus faecalis* and *Clostridium Cluster* IV. Except for *Faecalibacterium prausnitzii* and *Bifidobacteria*, samples in GT and MGIE reagents introduced comparable variations in *Lactobacillus* and *Clostridium Cluster* IV. Samples storage at ET_4 °C and in LS reagents caused dramatic increased abundance in all of the tested bacteria (Fig. 6 c).

### Do different sequencing platforms generate consistent results of identical samples?

Given that most of the microbiome results were obtained from different studies on different sequencing platforms, the consistency of microbiome data is largely overlooked in the context of different analyzing platforms. To test whether different sequencing platforms will introduce biases in microbiome data, replicated DNA samples extracted from same feces were shipped to two certified microbiome sequencing companies for sequence analysis on 16S rRNA gene according to their well-established protocols in our current study. The generated data were processed and analyzed by same researcher with same methods. The comparisons between the data from two sequencing platforms included α and β diversities, intestinal typing and community composition at phylum level. First, although our results indicated that the values of Shannon and Simpson indices from the two sequencing platforms were slightly varied under the same storage condition, the general change trends of α diversity under different storage conditions was consistent between the two platforms (Fig. 7a-b).

**FIG 7.**
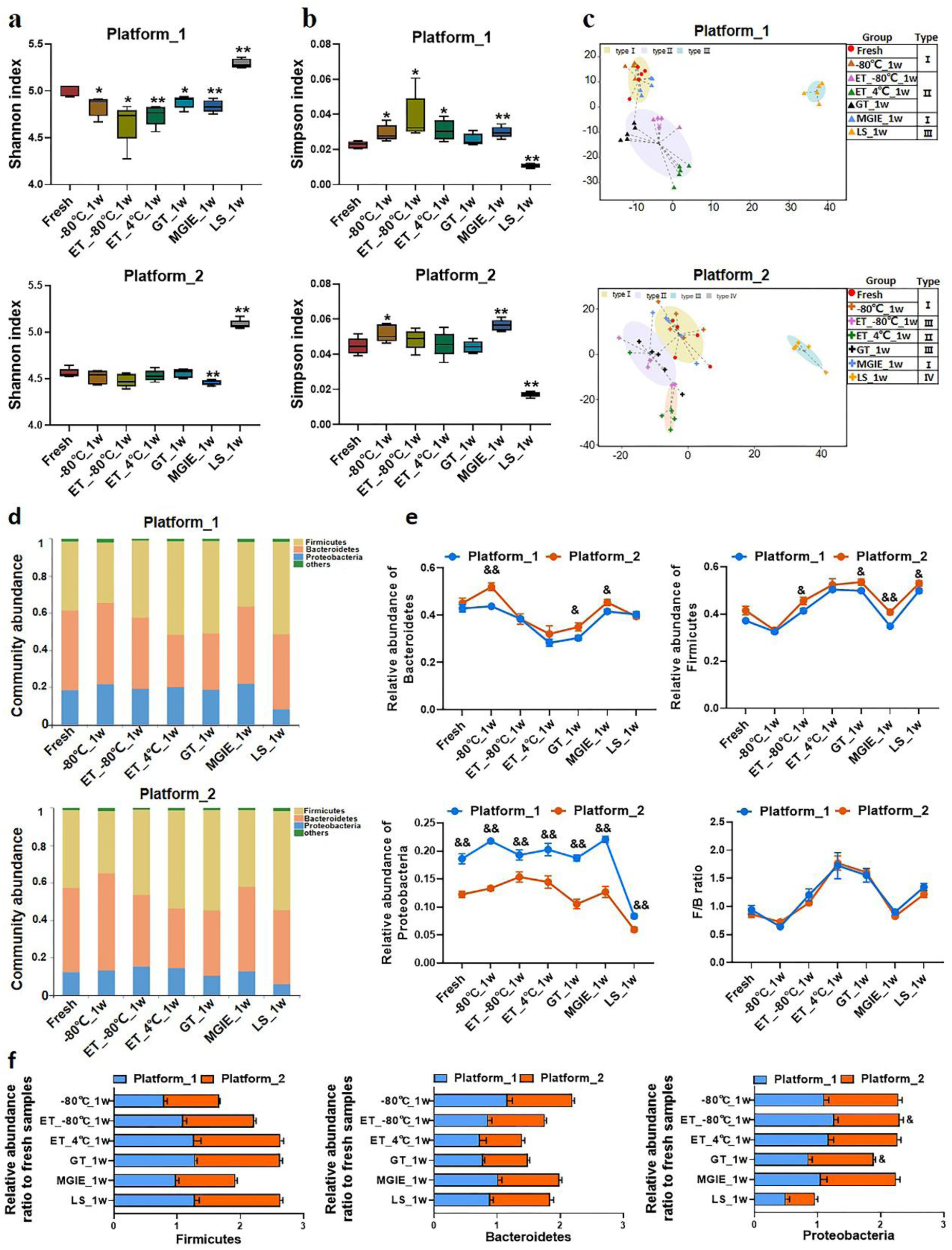
The impacts of different sequencing platforms on gut microbial profile. The community α diversity in platform_1 and platform_2 were analyzed by (a) Shannon diversity index and (b) Simpson diversity index. (c) The typing analysis on OTU level based on Jensen-Shannon Distance in platform_1 and platform_2. (d) Microbial communities under different storage conditions in platform 1 and platform_2. (e) The relative abundance of phyla Firmicutes, Bacteroidetes, Proteobacteria and the ratio of Firmicutes to Bacteroidetes under different platforms. (f) The relative abundance ratio (stabilized samples to fresh samples) of the dominant phyla in different platforms. * *p <* 0.05, compared with fresh group; & *p* < 0.05, && *p* < 0.01, compared with platform_1.

Then, the classification of dominant bacterial populations was performed with typing analysis based on Jensen-Shannon Distance. Samples from platform_1 were annotated into 3 types. Samples of fresh, −80°C and MGIE reagents were clustered type I, and samples of ET_-80 °C, ET_4 °C, and GT reagents in type II, except for those in LS reagents in type III. Although samples from platform_2 were annotated into 4 types, the cluster patterns were similar with platform_1, except for a sub-clustering within type II, in which ET_4 °C was differently classified with ET_-80 °C and GT reagents (Fig. 7c). The following community abundance analysis revealed the compositional structure of the dominant phyla between the same storage conditions including Firmicutes, Bacteroidetes, and Proteobacteria was comparable in data from the 2 sequencing platforms, even though the absolute values were of much difference, especially in Proteobacteria (Fig. 7d). Interestingly, similar to that of α diversity, the fluctuating trend of the dominant phyla under different storage conditions was consistent between the two platforms, which was clearly demonstrated by the overlapped F/B ratio under the 2 sequencing platforms (Fig. 7e). Finally, to assess the alteration of the dominant phyla introduced by sequencing platforms more accurately, we compared the relative abundance ratio (stabilized samples / fresh samples) of Firmicutes, Bacteroidetes, and Proteobacteria under the 2 sequencing platforms. No significant variation in the abundance of the dominant bacteria between two platforms, except for ratio of Proteobacteria in ET_-80°C and GT reagent (Fig. 7f).

Altogether, our results indicated that variations introduced by sequencing platform led to alterations of bacterial abundance, however, the trend of fluctuation is highly consistent under different storage conditions, but regardless of sequencing platforms. Thus, the variations introduced by storage conditions surpassed types of sequencing platforms.

## DISCUSSION

In the current study, we systemically evaluated the impacts of various fecal sample storage conditions, storage periods, and sequencing platforms on gut microbial profile based on 16S rRNA gene sequencing. Our results highlighted that gut microbiome profile varied to different extent under different sample storage conditions, which surpassed the impacts of storage periods and sequencing platforms.

There are huge amount of publications each year in the research on gut microbiota and human diseases (5, 34-36), however, data of gut microbiome usually exhibited dramatic variations and even inconsistency between studies (37). Generally, biases may be introduced during the whole experiment such as sample collection, transportation, storage, DNA extraction and so on (14, 38, 39). Commercial stabilizer for preserving fecal samples at room temperatures are usually proposed due to the fact of sampling without possibility for immediate freezing at −20 °C or even below and transportation in clinical study, especially in remote areas. Although the efficacy of some commercial stabilizers on maintaining the original status of gut microbiome has been compared, the uncertainty still exists because of the inconsistent observations among some reports (24, 26). Although previous studies suggested that inter-individual variation from donors superseded that introduced by storage conditions (25, 40, 41), the impacts of different storage conditions on microbiome profile need systemic evaluation. Therefore, in our current study, a multi-dimensional comparison among different storage conditions was performed by using a mixture of homogenized feces from rats to minimize the probable artifacts during sampling process. DNA extraction is also a critical step for introducing variation of microbiome data if different protocols or extraction kits were used(14). The genomic DNA of all samples was extracted by one experimenter with same protocol in our current study. Furthermore, many studies evaluated the effectiveness of preservation methods by using −80 °C or GT reagents stored samples as controls, whereas the efficacy of these preservation conditions are still inconclusive (20). Thus, a part of fecal sample divided from the homogenized mixture on-site was used as the fresh control for comparison with other storage conditions. Stabilization of fecal samples with ethanol is an easy and economical way, and therefore different concentrations of ethanol were used for fecal sample preservation in previous reports including 100% ethanol for spider monkey (24), 95% ethanol for human or dog (22), and 70% ethanol for canine fecal sample storage (20). However, inconsistent results were observed in different studies. For instance, Song *et al.* found that the preserving effect of 95% ethanol is comparable with FTA cards and OMNIgene Gut kit at ambient temperature, and they also strongly cautioned against the use of 70% ethanol (22), while Horng *et al.* reported that samples stored in 70% ethanol showed closest similarity with that of fresh samples, and therefor 70% ethanol was suggested as the best method for preserving canine fecal samples (20). These inconsistent conclusions might be associated with different fecal sample donors or different definitions for the “Fresh” samples, for example, Horng *et al.* processed canine feces for DNA extraction after 2 h of temperature treatment, whereas Song *et al.* extracted DNA of human and dog feces on the day of donation. Our current study concluded that −80 °C was suitable for rat samples preservation although some variations were observed, while storing samples at −80 °C with 70% ethanol showed advantages in long term storage. Notably, we found that samples at 4 °C with 70% ethanol showed low similarity to that of fresh samples, which is different from observations from Horng, K. R. *et al.*

Meanwhile, given its “excellent” performance in sample storage, GT reagent was usually considered to be an effective preservation methods and even used as control for evaluating the efficiency of other preservation methods (21, 22, 42). However, to the best of our knowledge, the similarity of microbiome profile between samples stored in GT reagents and fresh samples was unclear. Thus, in this study, we systemically evaluated the performance of three commonly used commercial stabilizers including GT, MGIE, and LS by comparing to fresh samples. Our results suggested that GT reagents might not be the most cost-effective reagents for fecal storage given its comparable performance in maintaining the original status of microbiome profile with other conditions such as MGIE, but relatively higher cost. On the contrary, the microbiome profile of samples in MGIE reagents was more similar with the fresh samples, whereas storage of samples in LS reagents introduced substantial variations in many aspects.

Given that gut microbiota plays critical roles in maintaining human health or disease development by producing microbial metabolites like bile acids or SCFAs (43, 44), the enrichment or depletion in the abundance of the functional bacteria that are SCFAs-producing or involved in bile acids metabolism is of great significance for investigating the microbial mechanism. However, very few attention has been paid to the variation that may be introduced during sample collection, storage condition or DNA extraction before sequencing up to now. In our current study, we specifically evaluated the abundance change of bacteria that are involved in SCFA-producing and/or bile acid metabolism using either 16S rRNA gene sequencing or qPCR. Our data suggested that different storage conditions could cause diversified fluctuations in the relative abundance of functional bacteria in a time-dependent or –independent way, and we also found that *Bacteroides* and *Lachnospiraceae_*NK4A136_group were relatively stable under these storage conditions. Moreover, the exact abundance of 6 functional bacteria in different storage conditions was qualified with samples stored for 6 months. We demonstrated that conditions of −80 °C, ET_ −80 °C, MGIE or GT reagents introduced comparably less variations, while conditions of ET_4 °C and LS reagents introduced higher variations, especially the striking increase of these bacteria in LS reagents. Consistently, the comparable storage effects between GT and MGIE were previously reported(21). It is the first time to visualize the impacts of different storage conditions on these functional bacteria quantitatively. Our current results highlight the importance of sample storage conditions, which may lead to dramatic biases in explaining the role of functional bacteria in disease, if different storage conditions were used.

Previous studies usually evaluated the impacts of short-term or long-term storage on gut microbial profile using samples stored ranging from 24h to 8 weeks (18, 40, 45). In our current study, the impacts of storage periods under different conditions were further evaluated in a multi-dimensional way after 1 or 2 weeks, and 6 months respectively. Our results showed that the dominant OTUs abundance of samples stored at −80 °C, ET_4 °C and MGIE reagents resulted in time-dependent alterations. Meanwhile, in comparison with fresh controls, samples stored at −80 °C, in either GT or LS reagents resulted in obvious variations in the dominant phyla time-dependently, while similar conclusion were not observed in the dominant genera. Actually, in most of the indicators, although some variations under different storage periods were observed, throughout the experiment, we found that variations between stabilized samples and fresh samples were larger than that of samples stored for different periods under the same storage conditions, indicating that variations caused by ways of storage condition were larger than that of storage period. Additionally, although variations in gut microbiome were mainly caused by the difference between individuals, sequencing is also a source for introducing variations in microbial data (25). Our current results showed, although α diversity and the relative abundance at phylum level varied between different platforms, the change trends between two sequencing platforms were still consistent among different storage conditions, indicating that biases introduced by sequencing platforms were less than that introduced by storage conditions.

## CONCLUSION

In the current study, we performed a multi-dimensional evaluation on the variations introduced by types of storage conditions, preservation period and sequencing platform on the basis of data acquired from 16S rRNA gene sequencing on rat fecal samples. Our results suggested that, compared to fresh control, the impacts on genomic DNA quality and yields are LS>GT>MGIE> ET_4 °C>-80 °C> ET_-80 °C in a time-independent way. Similarly, the impacts on α diversity are LS>-80 °C> ET_-80 °C>ET_4 °C> MGIE>GT in a time-independent way. The impacts on β diversity are LS>ET_4 °C>GT >-80 °C> ET_-80 °C> MGIE time-dependently. The impacts on abundance of functional bacteria are LS>ET_4 °C> ET_-80 °C>-80 °C>GT> MGIE. In addition, the impacts of storage conditions>storage periods>sequencing platforms. Therefore, our current results underpin that the storage conditions for fecal samples should be consistent to minimize the deviation that would influence the final readouts during the microbiome study, while same storage periods and protocols are also suggested.

## METHODS

### Animals

Male wistar rats were provided by Shanghai Slack Laboratory Animal Co., Ltd., The animal experiments were conducted under the Guidelines for Animal Experiment of Shanghai University of Traditional Chinese Medicine and the protocol was approved by the institutional Animal Ethics Committee.

### Commercial kits

Three commercial kits were used in the current study, including the Genotek OMNIgene·GUT OM-200 (DNA Genotek Inc., Canada), MGIEasy fecal sample collection kit (Shenzhen Huada Zhizao Technology Co., Ltd., China), and Longsee (Guangdong Nanxin Medical Technology Co., Ltd., China).

### Fecal sample collection and processing

Fecal samples from 10 rats were quickly collected into sterile 50 ml tubes and homogenized as much as possible. Next, these samples were aliquoted immediately, and aliquots were preserved using the following conditions: immediate freezing at −80°C with or without 70% ethanol (ET_-80°C), refrigerating at 4 °C with 70% ethanol (ET_4°C), the use of OMNIgene·GUT (GT), MGIEasy (MGIE) and Longsee (LS) according to the manufacturer’s instructions and stored for 1week, 2 weeks, and 6 months. Notably, according to the instructions, OMNIgene·GUT and MGIEasy samples were stored at room temperature, while Longsee samples were refrigerated at 4°C prior to DNA extraction. In addition to extracting DNA on the day of collection (fresh), extractions of other storage condition samples were conducted after 1week, 2weeks, and 6 months of storage, respectively.

### DNA extraction

DNA extraction was performed using QIAamp Power Fecal DNA Kit (QIAGEN, Germany). For samples immersed in solution, aliquots were centrifuged at 13,000 g for 5 min, and the supernatant was discarded. The pellet was washed with PBS, and centrifuged again at 13,000 g for 10 min. Subsequent DNA extraction steps were performed according to the manufacturer’s instructions. In addition, DNA concentration and the A260/280 ratio were tested by Colibri Microvolume Spectrometer (TIRERTEK BERTHOLD, Germany).

### 16S rRNA gene Sequencing

In our current study, DNA samples extracted at day 0, 1week, 2weeks, and 6 months of storage were applied to amplify the V3-V4 region of 16S rRNA gene using the universal primers 338F “ACTCCTACGGGAGGCAGCAG” and 806R “GGACTACHVGGGTWTCTAAT”. The sequencing was performed by the Illumina MiSeq PE300 system (Illumina, San Diego, USA) of Shanghai Meiji Biomedical Technology Co., Ltd., which is defined as platform_1 in the following experiments. Raw data files were demultiplexed, quality-filtered and merged by FLASH, and sequences whose overlap longer than 10 bp were merged using FLASH. The reads were clustered to OTUs with 97% similarity cutoff using UPARSE (version 7.1 http://drive5.com/uparse/) and chimeras were removed using UCHIME. The taxonomy of each sequence was analyzed by Ribosomal Database Project (RDP) Classifier algorithm (http://rdp.cme.msu.edu/) against the 16S rRNA gene database Silva (SSU123) using confidence threshold of 70%.

In addition, to investigate impacts of different sequencing platforms on gut microbial profile, identical samples extracted at 1week were analyzed using the Illumina Hiseq PE300 system (Illumina, USA) of BGI-shenzhen Co., Ltd, the V3-V4 region was amplified using the primers 341F “ACTCCTACGGGAGGCAGCAG” and 806R “GGACTACHVGGGTWTCTAAT”, which is defined as platform_2 in subsequent analysis. In order to obtain more accurate and reliable results in subsequent bioinformatics analysis, the raw data was filtered (46) and merged sequences whose overlap longer than 15 bp using FLASH (47). The tags were clustered to OUTs by scripts of software USEARCH (v7.0.1090) with a 97% threshold by using UPARSE (48), and chimeras were filtered out by using UCHIME (v4.2.40). OTU representative sequences were taxonomically classified using RDP Classifier v.2.2 against the database Greengene_2013_5_99, using confidence threshold of 0.6. Raw fastq files of this study were deposited in Sequence Read Archive database (accession number PRJNA561903).

### Quantitative RT-PCR

QRT–PCR was performed using SYBR Green (A25777, Thermo Fisher Scientific, USA), 96-well plates and the CFX connect Real-Time System. Each well was loaded with a total of 20 µl containing 2 μl of DNA, 0.5 μl of target primers, 7.5 μl of water and 10 μl of SYBR Select Master Mix. Hot-start PCR was performed for 40 cycles, with each cycle consisting of denaturation for 15 s at 94°C, annealing for 30 s at 60°C and elongation for 30 s at 72°C. Relative quantification was done using the 2-ΔΔCT method. Expression was normalized against universal primer of bacteria. Mean abundance of fresh samples were set as 100%. The primers used are shown in supplementary table S1.

### Statistical analysis

Alpha diversity was determined using Shannon’s diversity index, Simpson’s diversity index, Chao’s diversity index and Shannon’s evenness index, that calculated by mothur (version v.1.30.1). The Principal Coordinate Analysis (PCoA) based on weighted UniFrac, unweighted_unifrac, Bray_curtis and euclidean was conducted to evaluate similarity of microbial community. The classification of dominant bacterial populations under different storage conditions was studied mainly by typing analysis based on Jensen-Shannon Distance.

In our current study, statistical analyses were performed using the GraphPad PRISM version 8.0.1. Data obtained from experiments were shown as means±SEM, and differences between groups were assessed using two-tailed Student’s *t* test. Statistically significant differences were shown as follows: * *p* < 0.05, \*\**p* < 0.01, *** *p* < 0.001 compared with fresh group and & *p* < 0.05, && *p* < 0.01, &&& *p* < 0.001 compared with platform_1.

## SUPPLEMENTAL MATERIAL

**FIG S1** The effect of preservation methods on the community composition. (a) The ratio of Firmicutes to Bacteroidetes under different storage conditions. (b) Spearman correlation coefficients between the stored samples and corresponding freshly extracted ones on genus level. *p* < 0.05, \*\**p* < 0.01, compared with fresh group.

**TABLE S1** Primer sequences used in this study for microbial abundance.

## AUTHOR CONTRIBUTIONS

Junli Ma conducted the experiments, data analysis and manuscript writing; Lili Sheng revised the manuscript; Chuchu Xi helped in fecal sample collection and storage; Yu Gu and Ying Hong helped in data analysis; Ningning Zheng helped in uploading the data of 16s rRNA gene sequencing; Linlin Chen, Gaosong Wu, Yue Li, Juan Yan, Ruiting Han, Bingbing Li, Huihui Qiu and Jing Zhong helped in subsequent experiments. Wei Jia participated in the design of this study; Houkai Li supervised the project and revised the manuscript.

## ACKNOWLEDGMENTS

This work was funded by National Natural Science Foundation of China (No. 81873059 & 81673662), & National Key Research and Development Program of China (No. 2017YFC1700200), & Program for Professor of Special Appointment (Eastern Scholar) & Shuguang Scholar (16SG36) at Shanghai Institutions of Higher Learning from Shanghai Municipal Education Commission.

Program for Professor of Shuguang Scholar

